# Quantify genetic variants’ regulatory potential via a hybrid sequence-oriented model

**DOI:** 10.1101/2024.03.28.587115

**Authors:** Yu Wang, Nan Liang, Ge Gao

## Abstract

Understanding how noncoding DNA determines gene expression is critical for decoding the functional genome. Leveraging a hybrid sequence-oriented architecture, we developed SVEN to model (and predict) tissue-specific transcriptomic impacts for large-scale structural variants across over 200 tissues and cell lines. We expect that SVEN will enable more effective *in silico* analysis and interpretation of human genome-wide disease-related genetic variants.

## Main Text

Whole-genome sequencing enables high-resolution maps of genomic variations in human genome ^1^, revealing pervasive structural variants (SVs) in human genome^2^. Operationally defined as large-scale genomic alternation (>50bp), structural variants have been showed prominent impacts for several complex diseases, including schizophrenia, rheumatoid arthritis, and type 1 and type 2 diabetes^3^. Typically, SVs are thought to function via influencing gene expression by affecting regulatory regions of genes^4^.

Multiple efforts have been made for systematically characterizing the cellular impacts of genetic variants. In addition to the classical annotation-oriented approaches ^5^, multiple sequence-oriented methods have been proposed ^6,7^. Trying to “learn and model” regulatory codes from DNA sequences directly via various deep learning networks, sequence-oriented methods have demonstrated notable performance in predicting the expression influence for SNV and small indels, in both well-annotated and poor-annotation genomic regions ^8-10^.

SVEN employed a hybrid architecture to learn regulatory grammars and infer gene expression levels from promoter-proximal sequences in a tissue-specific manner (**Fig. 1a and Methods**). Briefly, we first trained multiple regulatory-specific neural networks based on 4,516 transcription factor (TF) binding, histone modification and DNA accessibility features across over 400 tissues and cell lines generated by ENCODE (**Extended Data Fig. 1**). Evaluations suggested that these networks successfully learned the underlying “regulatory codes” from inputs directly (**Fig. 1b**, also see **Extended Data Fig. 2**). A data-oriented feature selection procedure was further employed and excluded 802 networks associated with less-informative regulatory features based on model performance (**Extended Data Fig. 3 and 4**, and see **Methods** for more details). The output of remaining 3,714 networks, along with a separated mRNA-decay-related feature set ^11^, were feed to train 218 gradient boosting trees for inferring gene expression in 218 tissues and cell lines respectively.

**Fig. 1:**
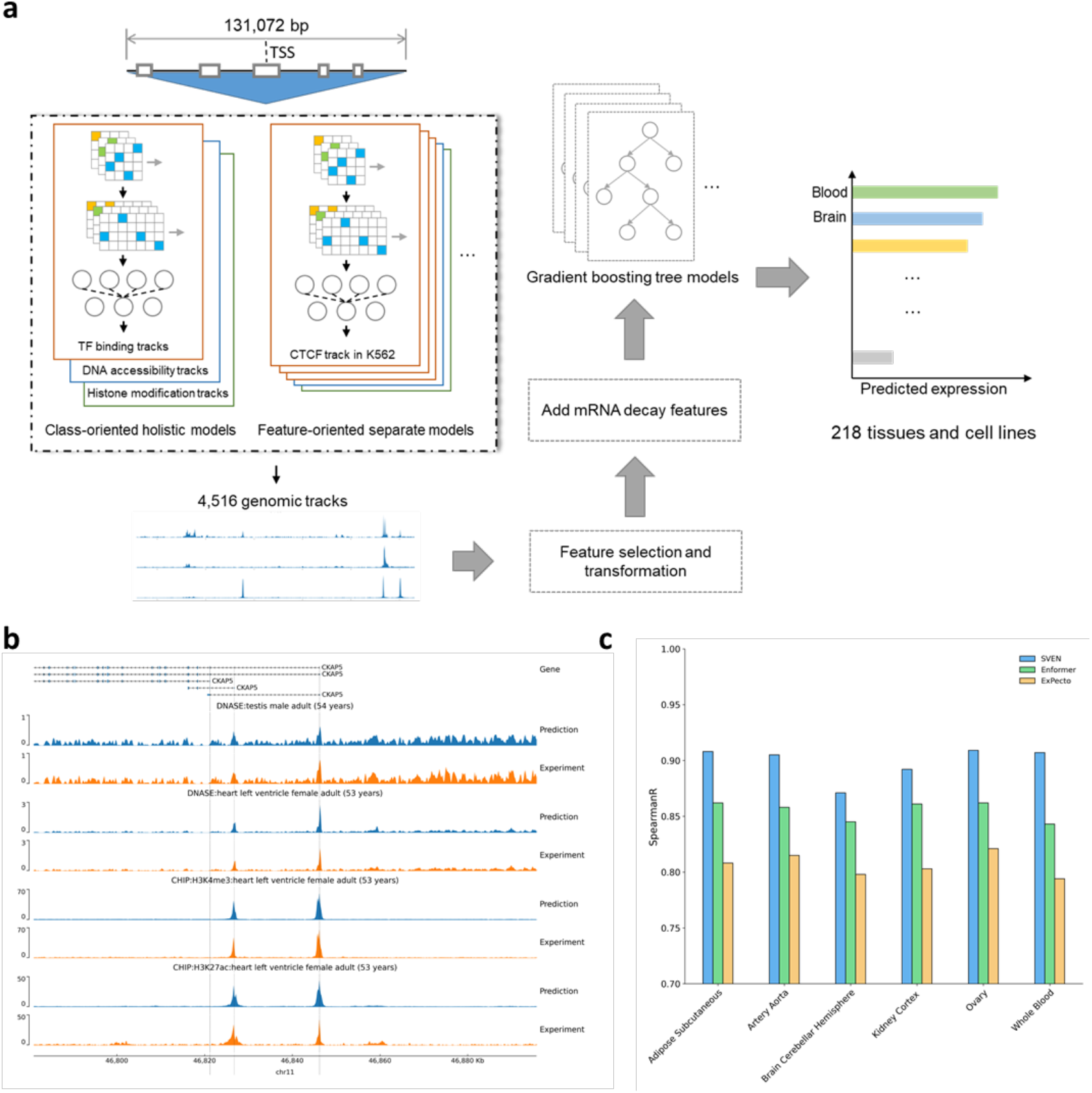
Tissue-specific gene expression prediction framework. **a**. Schematic overview of SVEN. SVEN consists of three components that act sequentially: the sequence-based deep neural networks to learn regulatory codes from sequences, feature selection and transformation to reduce the dimensionality of features, and gradient boosting tree models to predict gene expression level in tissue-specific manner. **b**. Representative example of observed and predicted functional genomic features (log_10_ scale) from deep neural networks. The sequence (chr11:46781020-46895708) at *CKAP5* locus is in the test dataset of networks. pyGenomeTracks ^25^ was used for plotting genomic tracks. **c**. SVEN outperform state-of-the-art tools in tissue-specific gene expression prediction on held-out testing sequences. Predicted log10RPKM were compared with RNA-seq observed log10RPKM in 6 example tissues that reported by Enformer and ExPecto. Spearman correlations between predicted and observed values were calculated for comparison.

To evaluate model performance in gene expression prediction, we assessed SVEN on held-out testing sequences. SVEN made accurate predictions of gene expression levels across 218 tissues and cell types, with mean Spearman correlation 0.890 (**Supplementary Table 1**). Of note, SVEN showed consistently better performance across different tissues than ExPecto ^8^ and Enformer (**Fig. 1c** and **Extended Data Fig. 5**). We then evaluate SVEN’s performance in predicting SVs’ regulatory impact by applying SVEN to a large-scale SV dataset from 32 diverse human genomes with paired RNA-seq data ^12^ (**Methods**). Out of 114,904 SV-gene pairs in 26 samples (corresponding to 51,335 SVs and 20,318 genes, **Extended Data Fig. 5**), SVEN accurately predicted their regulatory effects with Spearman correlation 0.910 between predicted and observed expression level from paired RNA-seq data (**Fig. 2a**).

**Fig. 2:**
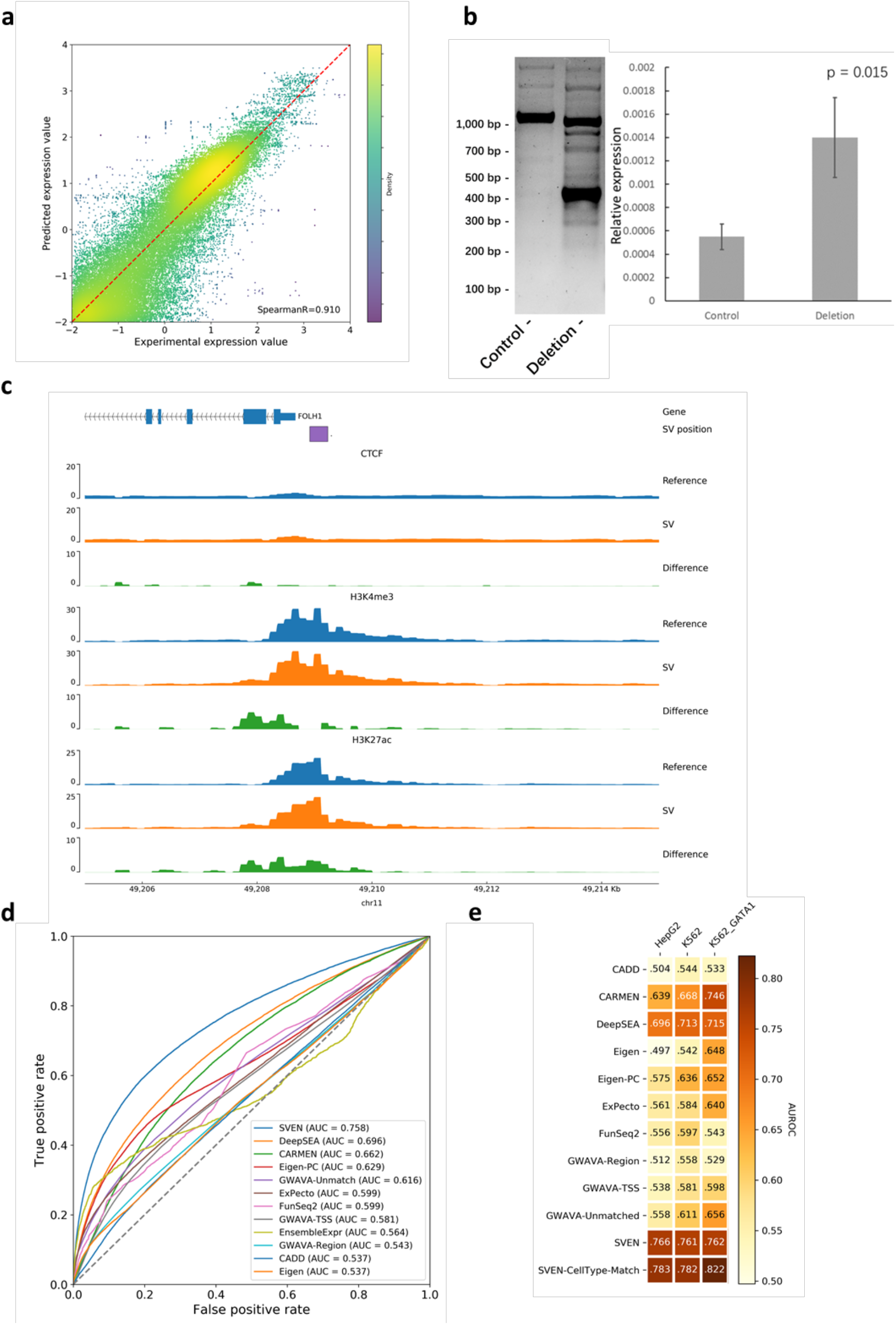
SVEN can quantify genetic variants’ regulatory potential accurately. **a**. Evaluation of SVEN in assessing the effects of SVs on gene expression. Predicted log_10_RPKM from GM12878 model was compared with observed RNA-seq log_10_RPKM in LCL cell line. Non-expressed genes were removed, Spearman correlation was shown and color represents the density of dot. **b**. (left) DNA fragments generated by PCR on genomic DNA from control A375 cells and those with CRISPR deletion on the FOLH1-SV region. (right) The effect of CRISPR deletion on the relative abundance of mRNA for *FOLH1* as determined by real-time quantitative PCR analysis. The reference gene was *RPL41*. Since there was difference between target deletion and designed deletion in CRISPR experiment, we also used SVEN to predict the expected deletion in CRISPR experiment based on the position of sgRNA, and it came to same conclusion. **c**. Annotations of FOLH1-SV by SVEN annotation model. Difference annotation tracks shown difference between the signal of reference sequence and the signal of deleted sequence (*Signal*_*d*_ *– Signal*_*r*_). Since the SVEN annotation did not include annotation in A375 cell line, the CTCF, H3K4me3 and H3K27ac signal shown here were features with the highest feature contribution in SVEN model (A375). **d**. The ROC curves on small noncoding variants measured by *in vitro* massively parallel reporter assays. We use 5,248,124 variants tested in a variety of cell types, including 30,121 positive variants and 5,218,003 negative variants ^18^. For each variant, we used the predicted difference of 4,516 annotations between the reference allele and alternative allele and generated a sum of the difference without any additional training to evaluate variants’ effects. **e**. Model’s performance on cell-line-specific variants. To evaluate the performance on cell-line-specific variants, we extracted variants tested in HepG2 and K562 cell-line from the benchmarking dataset, selecting 265 and 483 cell-line matched features in HepG2 and K562 cell-line from 4,516 SVEN annotations respectively. SVEN-CellType-Match used same K562 annotations for K562 and K562_GATA1 variants.

To further demonstrate the predictive ability of SVEN, we selected deletions from the large-scale SV datasets for experimental validation (**Supplementary Table 2,**and see **Methods** for more details). For top 5 deletions with the highest regulatory impacts predicted by SVEN (**Supplementary Table 3**), we ran independent CRISPR-based assays. The results showed that SVEN predicted the direction of the impact of 4 deletions on gene expression successfully, and 2 of them were statistically significant (**Fig. 2b, Extended Data Fig. 6**). Of note, while the deletion at the upstream of cancer biomarker PSMA-encoding gene *FOLH1* ^13^ disrupts the promoter region (**Extended Data Fig. 6**) and the annotation-based algorithm predicted that it barely affect gene transcription (regulatory disruption score=-0.02) ^14^, SVEN gave an increment prediction correctly, partly due to its annotation module indicated that variant effectively increases the expression-active H3K4me3 and H3K27ac signal rather than the deletion of known silencer^15^ or insulator, which further suggest plausible underlying mechanisms for this deletion (**Fig. 2c**).

In attempts to estimate SV’s regulatory effects genome-widely, we systematically screened a in large-scale dataset derived from 3,622 samples ^16^. For 165,607 curated SV-gene pairs corresponding to 73,829 SVs (**Methods**), SVEN suggested most of SVs did not affect gene expression significantly (**Extended Data Fig. 7**). On the other hand, we noticed that known pathogenic deletions are more likely to affect gene expression, especially exon-disrupted SVs with longer length (**Extended Data Fig. 7**). Interestingly, SVEN suggests these pathogenic SVs tended to decrease gene expression and the loss of function of target genes might be one of reasons of their pathogeny. For example, the SV nssv17171470 (chrX:139530701-139530862) is a 161 bp deletion located at the locus of the gene *F9*, which encodes coagulation factor IX and is associated with hemophilia B. SVEN predicted that it would decrease the expression of *F9* (log2 fold change = -10.83 in liver). This deletion disrupts the first exon and the promoter region of *F9* (**Extended Data Fig. 7**), including the TF binding sites of HNF4A that has been reported to have positive and significant correlation with the level of F9 in liver ^17^.

As a sequence-oriented model, SVEN should also work with small noncoding variants. Therefore, we further evaluated SVEN’s performance on variants whose functional effects were directly measured by *in vitro* massively parallel reporter assays (MPRAs) ^18^. Compared with several dedicated state-of-the-arts, SVEN showed the best performance with AUROC improved by 8.9% (**Fig. 2d**). Of note, SVEN cell-line matched prediction outperformed cell-line agnostic predictions (**Fig. 2e**), further demonstrating the importance of SVEN’s context-specific design.

While feature analysis showed that TF binding makes high contribution (**Methods and Extended Data Fig. 8**), the DNA accessibility features showed the lowest mean importance across all 218 models, which may partly due to their bidirectional biochemical nature: an accessible region of DNA can be bound by either activating or repressing regulatory factors. Of note, we found that mRNA decay features had the highest mean feature importance (**Extended Data Fig. 8**), likely owing to the close relationship between mRNA decay rate and the regulation of gene expression ^19^. We used decay constants to control the receptive field of transformed features (**Methods and Extended Data Fig. 3**), and the preference of different category of features was diverse. For DNA accessibility features, those with the largest receptive field had the highest feature importance, suggesting a non-neglected role for distal regulation (**Extended Data Fig. 8**). Meanwhile, the feature importance of histone modification features shown similar pattern, but the difference was the features from TSS-proximal regions shows the highest importance, further confirming the importance of histone modifications ^20^ (**Extended Data Fig. 8**).

We believe that SVEN, as an accurate and flexible sequence-oriented model, will enable more effective and efficient mining disease-related genetic variants in human genome. Thus, we have made the whole package of SVEN, along with tutorials and demo cases, available online at https://github.com/gao-lab/SVEN for the community.

## Supporting information

Extended Figures as well as the Table Legends of Supplementary Table 1-4

## Data availability

Basenji2 training, validation and testing sets as well as trained model were obtained from https://console.cloud.google.com/storage/browser/basenji_barnyard. ExPecto training, testing sequences on tissue-specific gene expression prediction as well as TSS annotation of genes was obtained from https://github.com/FunctionLab/ExPecto. Trained Enformer was obtained from https://github.com/deepmind/deepmind-research/tree/master/enformer. The performance of ExPecto and Enformer on RNA-seq data in six example tissues was obtained from their published papers. SVs as well as paired RNA-seq data from 26 samples were obtained from https://www.internationalgenome.org/data-portal/data-collection/hgsvc2. SVs from 3,622 Icelanders were obtained from https://github.com/DecodeGenetics/LRS_SV_sets. Pathogenic SVs were obtained from dbVar at https://www.ncbi.nlm.nih.gov/dbvar/studies/nstd102/. Gene annotation was obtained from https://www.gencodegenes.org/ (v24, GRCh38).

## Code availability

All codes for training and evaluating models, as well as trained models and detailed paraments are available at https://github.com/gao-lab/SVEN.

## Author Contributions

G.G. conceived the study and supervised the research. W.Y. designed and implemented the computational framework and conducted benchmarks and case studies with guidance from G.G.

L.N. conducted experimental validation. W.Y. and G.G. wrote the manuscript with inputs from all authors.

## Acknowledgements

This work was supported by funds from the National Science and Technology Major Project (grant no. 2022ZD0115004), the Changping Laboratory, as well as the State Key Laboratory of Protein and Plant Gene Research and the Beijing Advanced Innovation Center for Genomics at Peking University. Part of the analysis was carried out on the Computing Platform of the Center for Life Sciences of Peking University and supported by the High-performance Computing Platform of Peking University and Changping Laboratory.

