## Extended Figures as well as the Table Legends of Supplementary Table 1-4 for "Quantify genetic variants’ regulatory potential via a hybrid sequence-oriented model"

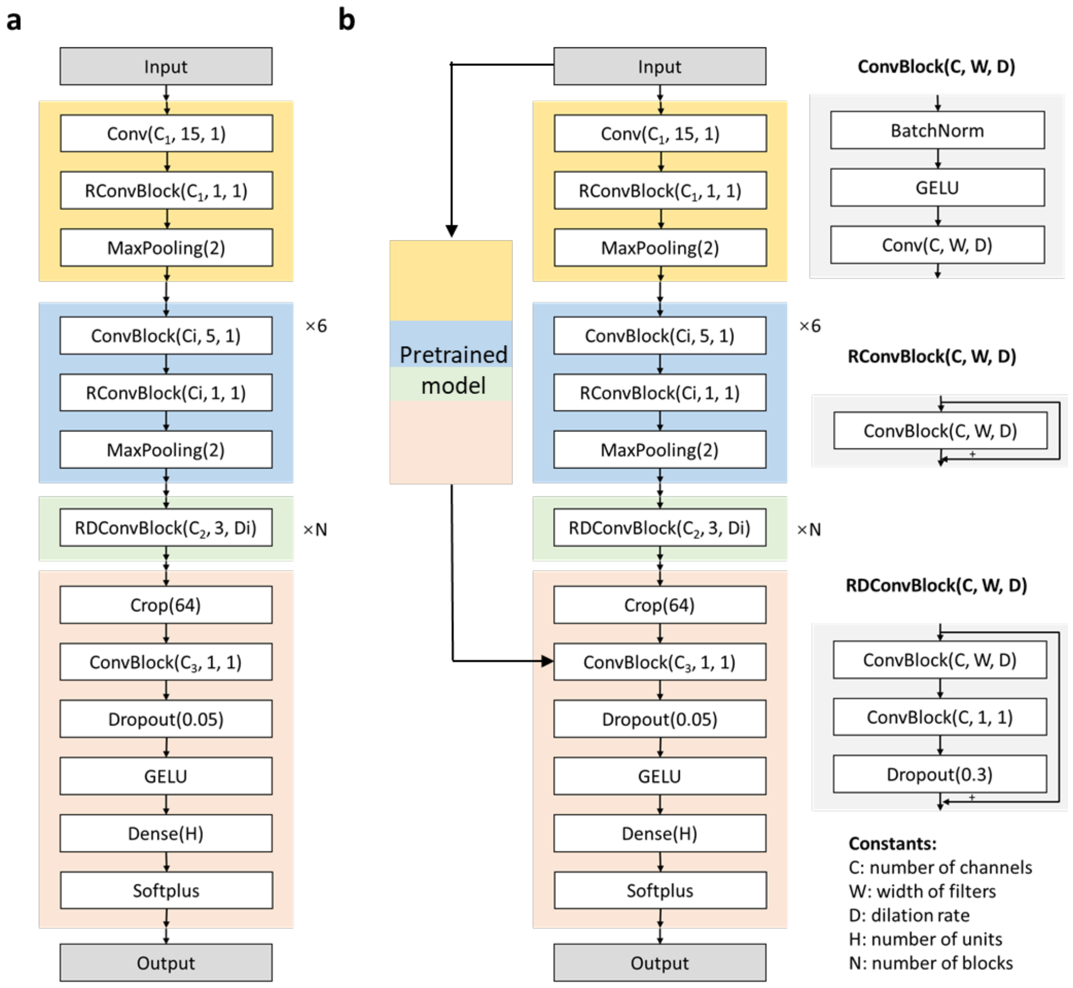


**Extended Figure 1: Model architecture of deep neural networks. a.** Basic model structure of deep neural networks. **b.** Model structure for histone modification and TF binding models.

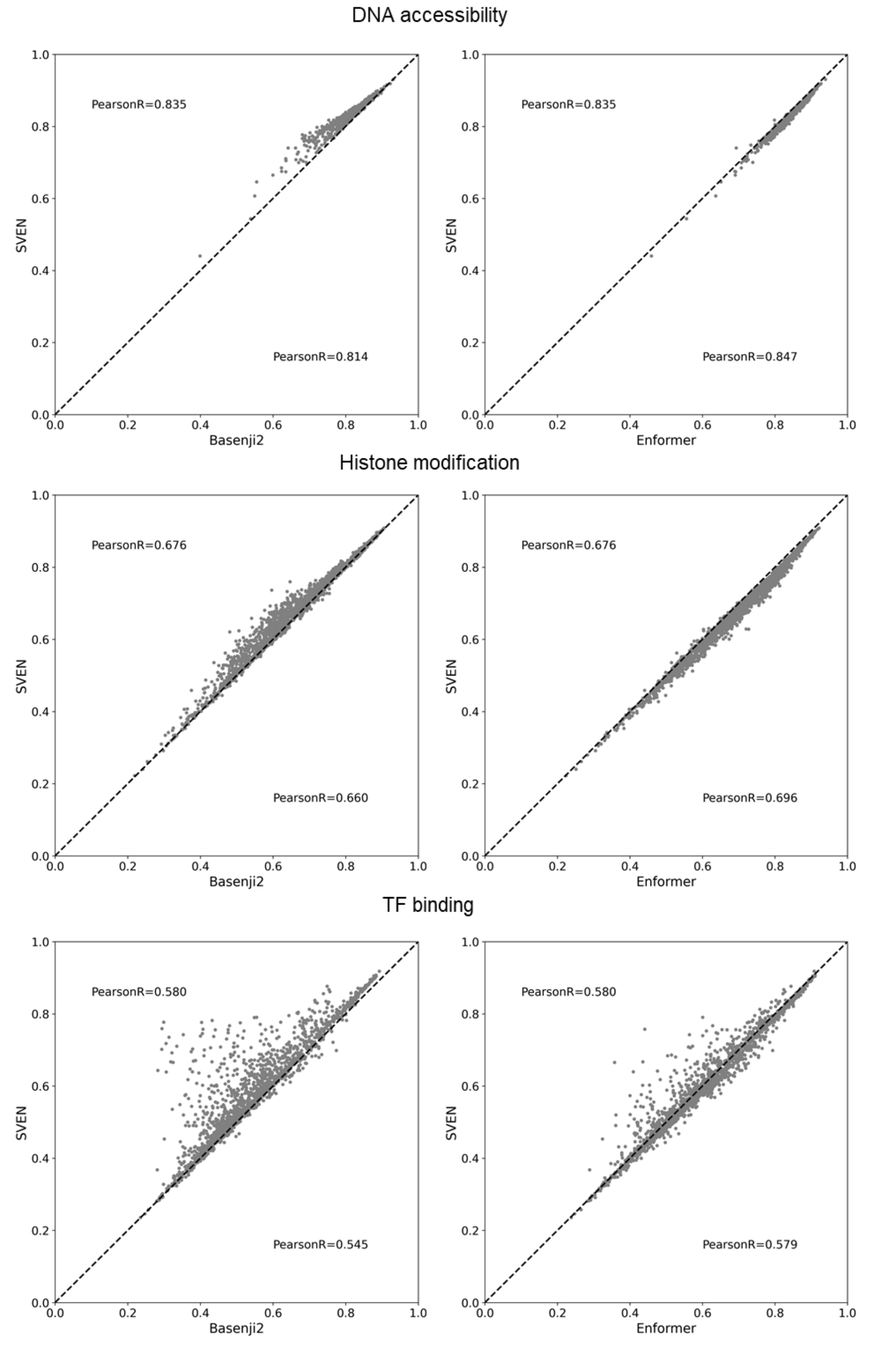


**Extended Figure 2: Model performance comparison in predicting functional genomic signals.** We compared the performance of deep neural networks of SVEN with state-of-the-art tools (Basenji2 and Enformer) in 4,516 DNA accessibility, histone modification and TF binding features. Pearson correlations are shown. Compared with Basenji2, we showed better performance in genomic signal prediction, and achieved similar performance with Enformer, which takes 50% longer sequence as the input.

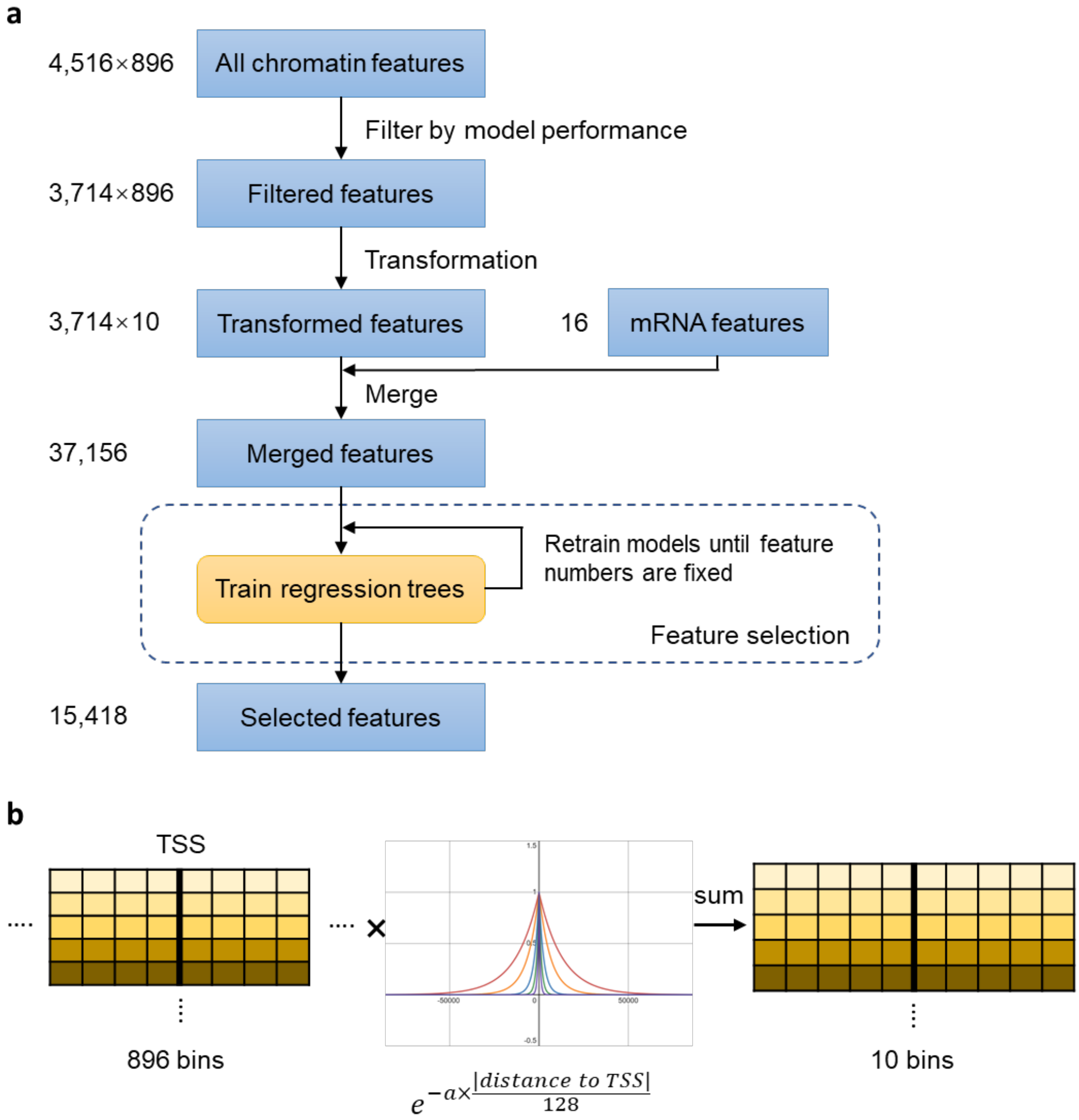


**Extended Figure 3: Schematic overview of feature selection and transformation. a.** Schematic overview of feature selection procedure. We first filtered genomic features according to model performance (Pearson R≥0.5). Then we transformed features and used transformed features and mRNA decay features to train feature selection models until the model performance decreased or the number of features no longer dropped. **b.** Illustration of feature transformation. We used 10 exponential functions with 5 decay constants (*a* in equation) to transform features.

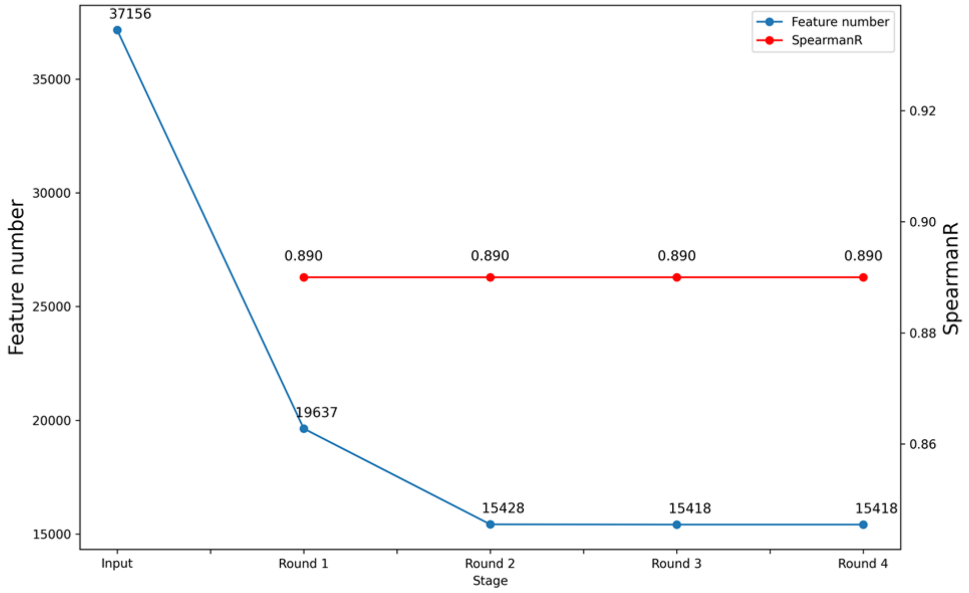


**Extended Figure 4: Mean feature number of all 218 models in feature selection procedure.** Input features are transformed filtered features and mRNA decay features. Mean feature number and Spearman correlation are shown.

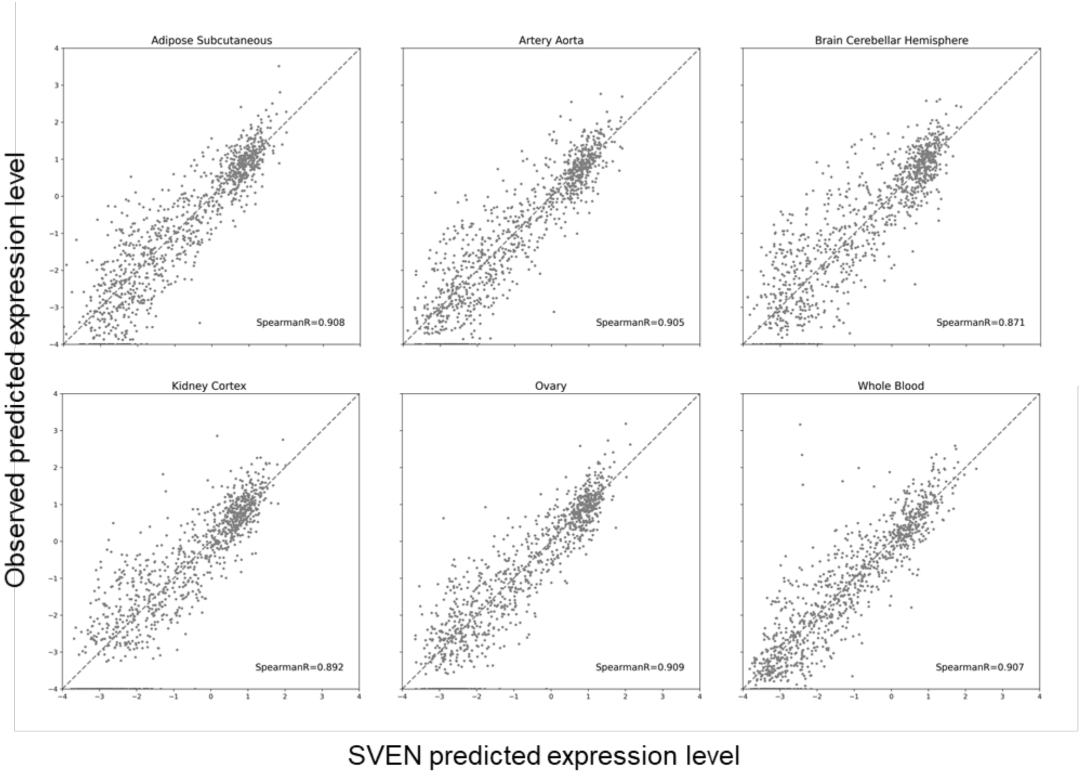


**Extended Figure 5: SVEN's tissue-specific gene expression prediction.** Predicted value by SVEN versus observed value for all testing sequences in 6 example tissues.

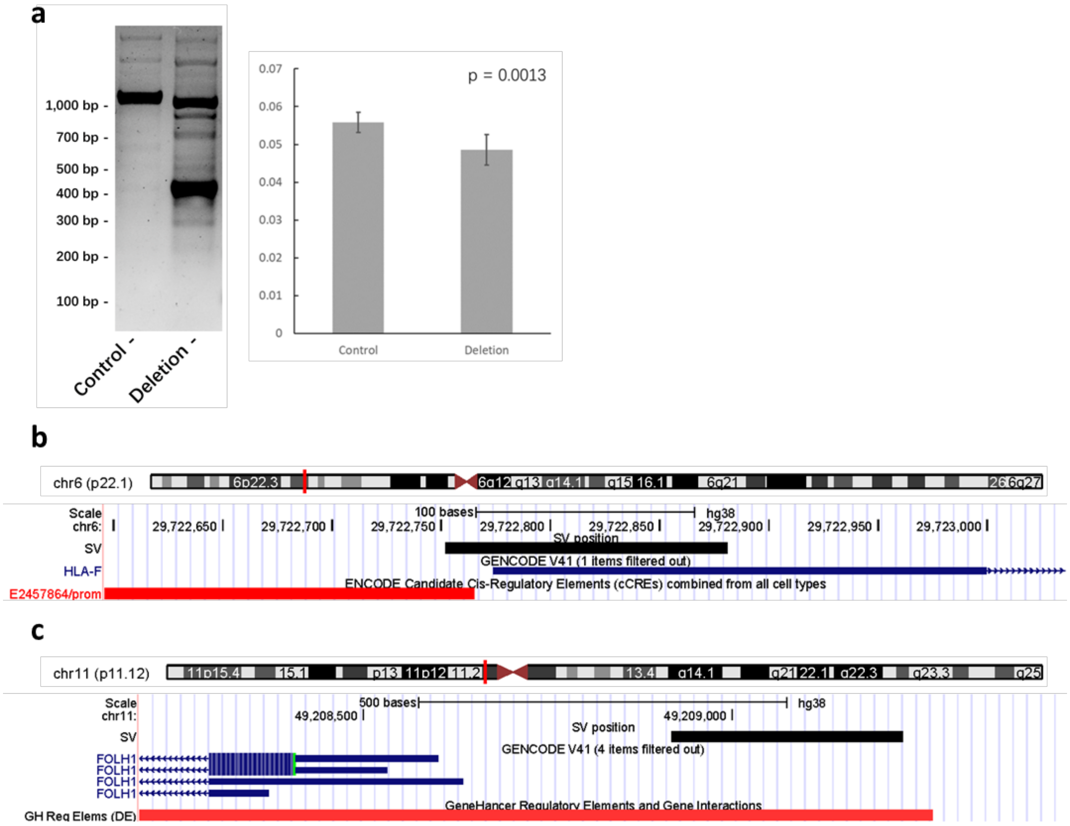


**Extended Figure 6: Experimental validation of HLA-F-SV and visualization of the HLA-A-SV and FOLH1-SV in UCSC genome browser. a.** (left) DNA fragments generated by PCR on genomic DNA from control A375 cells and those with CRISPR deletion on the HLA-F-SV region. (right) The effect of CRISPR deletion on the relative abundance of mRNA for *HLA-F* as determined by real-time quantitative PCR analysis. The reference gene was *RPL41*. **b.** HLA-F-SV (chr6:29722752-29722881) affects the promoter region and the first exon of gene *HLA-F*. E2457864 is promoter-like signature. **c.** FOLH1-SV (chr11:49208919-49209232) affects the promoter region (in red) of gene *FOLH1*. The annotation of promoter is from GeneHancer.

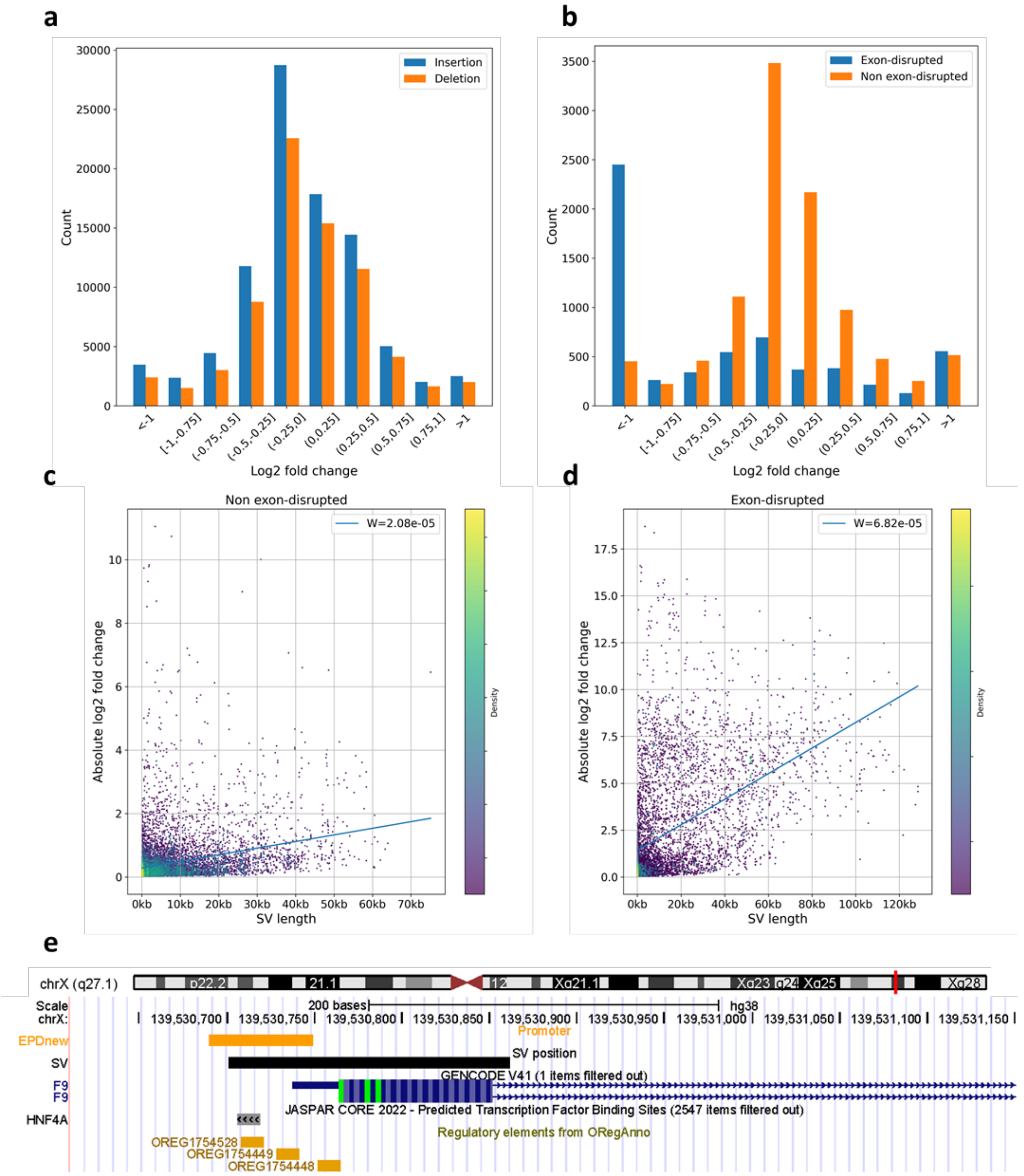


**Extended Figure 7: Effects of SVs on gene expression. a.** Effects of SVs from large-scale population on gene expression in 53 GTEx tissues. We shown the max log2 fold change (max of absolute value) of SVs across 53 tissues. **b.** Effects of pathogenic SVs on gene expression. **c and d.** Length of exon-disrupted (c) and non-exon-disrupted pathogenic SVs (d) versus effects on gene expression (absolute log2 fold change). Slopes (*W*) of fitted linear regression models are shown in upper right. Color represents the density of dot. **e.** SV nssv17171470 (chrX:139530701-139530862) disrupts the promoter and the first exon of gene *F9*. The ORegAnno IDs OREG1754528, OREG1754449, and OREG1754448 are transcription factor binding sites of HNF4A. The annotation of promoter is from EPDnew.

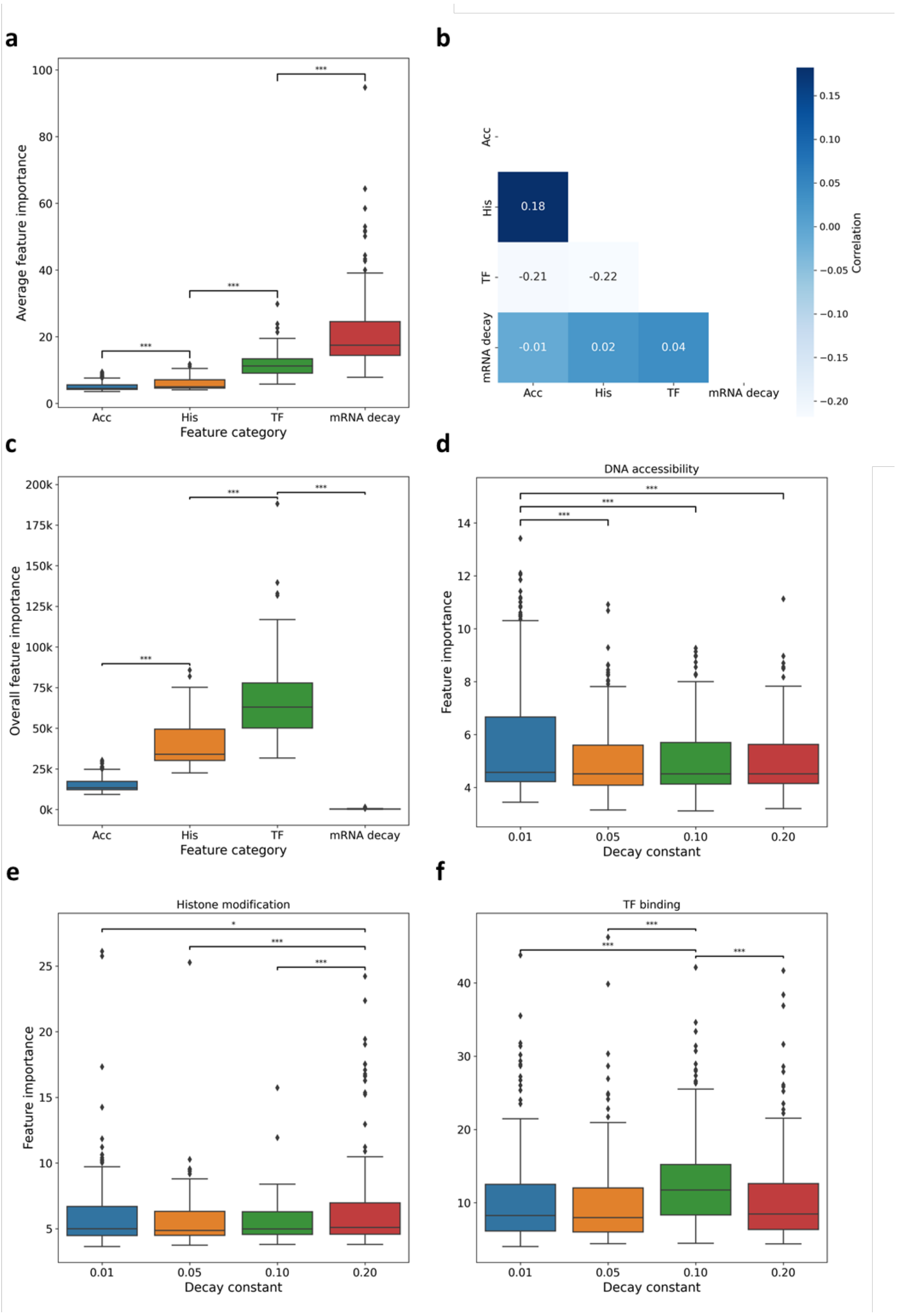


**Extended Figure 8: Feature importance for the prediction of gene expression. a.** Feature importance across all 218 models by feature category. Mean feature importance was calculated in each model. *: $p<0.1$; **: $p<0.01$; ***: $p<0.001$, same in (c), (d), (e) and (f). **b.** Feature contribution correlation between different categories across all models. Mean SHAP value was calculated in each model on all training and testing sequences. Spearman correlation are shown. **c.** Overall feature importance across all 218 models by feature category. **d, e, and f.** Feature importance of DNA accessibility (d), histone modification (e) and TF binding (f) features by decay constant used in feature transformation. We used different decay constant to control the receptive field of transformed features (**Methods**). 0.01~TSS ± 60 kb; 0.05~ TSS ± 20 kb; 0.10~ TSS ± 10 kb; 0.20~ TSS ± 5 kb.

### Supplementary Tables

**Supplementary Table 1** SVEN's performance of gene expression prediction on 218 tissues and cell lines.

**Supplementary Table 2** Criterion of selecting deletions for experimental validation.

**Supplementary Table 3** Top 5 deletions with the highest regulatory impacts predicted by SVEN from large-scale SV datasets for experimental validation.

**Supplementary Table 4** Guide RNA and PCR primers used in experimental validation.
