## Supplementary material for "Quantify genetic variants’ regulatory potential via a hybrid sequence-oriented model": Online Methods

**Framework architecture of SVEN**

The SVEN framework consists of three components that act sequentially. The first component is sequence-based deep neural networks to learn regulatory codes from sequences to predict functional genomic features like TF binding, histone modification, and DNA accessibility. Then we applied feature selection and transformation to reduce the dimensionality of generated features from deep neural networks. And finally, we used selected features as well as mRNA decay features to train gradient boosting tree models and make tissue-specific gene expression prediction.

The first component is deep neural networks to predict 4,516 functional genomic features including 1,896 TF binding features, 1,976 histone modification features, and 684 DNA accessibility features. Inspired by previous work [9], the basic neural network consists of three parts: (1) 7 residual convolutional blocks with pooling layers, (2) 11 residual dilated convolutional blocks, and (3) a cropping layer followed by pointwise convolution block and fully connected layer for the output. The input sequence of models is one-hot-encoded DNA sequence of 131,072 bp and the output is the (predicted) functional genomic features with length of 896 corresponding to centered 114,688 bp aggregated into 128-bp bins. The residual convolution blocks with pooling layers were used to extract sequence motifs and learn the interactions between them, and the dimension was reduced to 1,024. Then we used residual dilated convolutional blocks to learn long-range interactions across sequences. Finally, we applied a cropping layer to trim off 64 units at the beginning and end of the sequence due to the potential loss of information of these regions, where can only observe one-side information. Then we used pointwise convolution block to change the number of channels and following a fully connected layer with soft plus function as the activate function as the final output layer.

Inspired by our previous work [21], we introduced hybrid architecture to train deep neural network: class-oriented holistic models and feature-oriented separated models (**Extended Data Fig. 1**).

Specifically, we trained 3 class-oriented holistic models for each type of chromatin profiles (TF binding, histone modification and DNA accessibility), which could benefit from related features (such as same binding protein or modification across different cell lines and tissues). We also trained feature-oriented separated model, which paid more attention on sequence information alone, and each model corresponding to one feature. And finally, we selected the best models according to their performance on validation set and combine the outputs of these models as final output.

Besides, we found that DNA accessibility information could improve the model performance of TF binding and histone modification models, therefore, we incorporated pretrained DNA accessibility model into TF binding and histone modification models (**Extended Data Fig. 1**). Specifically, we first trained DNA accessibility holistic model as pretrained model. Then we incorporated pretrained model into TF binding and histone modification model, and the pretrained model was not frozen and was involved in further model training. The pretrained model was trained on the same sequences with TF binding and histone modification models.

The second component is feature selection and transformation. First, we filtered all 4,516 genomic features according to the model performance (**Extended Data Fig. 3**). We removed 802 features with Pearson $R < 0.5$ and left 3,714 genomic features. Then we transformed these features by the method described in ExPecto [8]. Briefly, we transformed features with 10 exponential functions to weight the upstream and downstream regions of TSS separately based on the assumption that regions with longer distance to TSS usually has less effects on gene expression, to reduce the dimensionality of features. We also use 5 different decay constants {0.01, 0.02, 0.05, 0.10, 0.20} to control the receptive field of transformed features. The number of features decreased from 3,327,744 (3,714×896) to 37,140 (3,714×10).

Besides, we also incorporated mRNA decay features for following model training. We extracted the GC content and the length of 3'UTRs and 5'UTRs from ensembl (104, GRCh38). We calculate the min, max, median, and mean value of all 3'UTRs and 5'UTRs for the target gene and generated 16 features for each gene. If there was no known 3'UTR or 5'UTR for the target gene, the value of feature was set to 0.

We combined the 37,140 transformed chromatin features and 16 mRNA decay features for further feature selection. We used all 37,156 features to train 218 extreme gradient boosting (XGBoost) regression tree models and each model corresponding to one tissue or cell line. Then we selected features used in trained models and then retrained models with selected features and repeated this procedure for several times until the model performance decreased or the number of features no longer dropped. We found that the number of features of all models no longer decreased after the third round of selection (**Extended Data Fig. 4**). Therefore, we used the features after the third-round selection as the final feature set and the mean feature number of all models was 15,418. And we also found that only features transformed by 4 decay constants {0.01, 0.05, 0.10, 0.20} were selected.

The third component is 218 gradient boosting tree models and each one corresponding to one tissue or cell type. We used selected features to retrain all XGBoost regression tree models as final models.

**Model training and evaluation of SVEN**

The first component of SVEN, the deep neural networks were trained, evaluated, and tested on the same sequences of 4,516 ENCODE (The Encyclopedia of DNA Elements) chromatin features extracted from 5,313 features used in Basenji2. The dataset contains 34,021 training, 2,213 validation, and 1,937 test sequences. The length of sequence was 131,072 bp and all sequences were based on GRCh38 reference genome.

We used Poisson negative log-likelihood function as loss function and the Adam as optimizer with default parameters. All models were implemented on TensorFlow (v2.5.0), and were trained on 8 NVIDIA Tesla A100 GPUs with batch size of 32 for 1,000 epochs with early stopping. The validation set was used for hyperparameters selection and model selection, and the performance on test set was reported as the final performance of all models. Pearson correlation was used to evaluate model performance, identical to the metric used by Basenji2 and Enformer. We used pretrained Basenji2 and Enformer model for model performance comparisons.

For the third component of SVEN as well as feature selection, we used same XGBoost regression tree models. We used same training and evaluation dataset as well as representative TSS of genes (lifted to GRCh38 coordinates) used by ExPecto. Briefly, the expression profiles of 218 tissues and cell lines were from GTEx, Roadmap and ENCODE project. We only used expression profiles of protein coding genes (18,632) and lincRNA genes (5,068). Then we added a pseudocount 0.01 (0.0001 for GTEx tissues due to high coverage) and applied log 10 transformation for model training. We also used trained deep neural networks to annotated all sequences centered by the TSS of each gene on all genomic signals for following feature selection and model training.

All genes from chromosome 8 were held-out for evaluation (987 genes) and all other genes were used for model training (22,713 genes). However, in order to search hyperparameters of regression tree models, we further selected all genes from chromosome 9 (939 genes) from training set as temporary validation set and the rest of genes (21,744 genes) were used for training models. Then we evaluated the performance of model on the temporary validation set for hyperparameters selection. After hyperparameters selection, we used fixed hyperparameters to retrain models for feature selection with whole training dataset (22,713 genes) and evaluated model performance on held-out test set as the model performance. All models were trained on NVIDIA Tesla A100 GPUs. Spearman correlation was used to evaluate model performance, identical to the metric used by ExPecto and Enformer. We used reported performance by ExPecto and Enformer for model comparisons.

**Feature importance analysis**

We used the built-in function of XGBoost to get the feature importance in trained regression tree models. The importance type we used was “gain” (the average gain of splits which use the feature). To investigate the contribution of features to the prediction, we calculated the SHAP [22] value of each feature on all training and testing sequences. Then we calculated the mean feature importance of each category or decay constant in each model and combined the feature importance or feature contributions from all 218 models for comparison. Single-tailed Wilcoxon-test and Spearman correlation was used in statistical analysis.

**SV expression effect prediction**

The effects of SVs on gene expression were estimated by the differences between the predicted expression level of sequences with reference allele and alternative allele. Here, we used log2 fold change to measure the effects of SVs on target genes. First, we checked the reference allele and alternative allele of SVs. The sequence information of SVs is necessary for SVEN, which is a sequence-based framework. Then, we checked the position of SV that whether the whole SV (all bases) fell in the region of 131,072 bp centered with TSS of any genes. If there was no overlapped gene, we would exclude this SV from further expression effect prediction. The TSSs of genes were same with the representative TSSs used in model training. For involved genes, we constructed SV-gene pairs and predicted the effects of SV on all involved genes. Specifically, we extracted the sequences of each gene from 5' to 3' and generated reference sequences and alternative sequences. If no reference allele was specified, we would use the sequences form reference genome (GRCh38) as reference sequences directly. Then we used SVEN to predicted the expression level of involved genes with reference and alternative sequences respectively. As the output of SVEN was log transformed value of expression level, we transformed predicted values to original values and then calculated the log2 fold change of gene expression level of involved genes as final output.

**Evaluation of SVEN's SV expression effect prediction**

To evaluate SVEN's prediction of SVs' effects on tissue-specific gene expression, we applied SVEN on a large-scale SV dataset from 32 diverse human genomes generated by Ebert *et al.* [12] using long-read sequencing. The dataset contains 107,578 SVs, including 66,189 insertions and 41,389 deletions. We first filtered insertions and deletions falling into the 131 kb sequences centered with the TSS of lincRNA and protein coding genes and constructed SV-gene pairs. We got 145,626 SV-gene pairs corresponding to 64,704 SVs and 21,448 genes.

We also downloaded paired RNA-seq (single-end) fastq files from Human Genome Structural

Variation Consortium Phase 2 (HGSVC2) and got paired RNA-seq data for 26 samples. RNA-seq data were mapped with HISAT2 (v2.2.1) [23] to reference genome GRCh38. The hisat2-index was built with reference genome GRCh38 and transcripts information extracted from GENCODE (v24) annotation. Then we mapped RNA-seq reads using hisat2 with default parameters. The gene abundance was generated by StringTie (v2.2.1) [24]. We used expression estimation mode to estimate the coverage of the transcripts on the basis of GENCODE (v24) annotation. The value of FPKM was calculated as the mean value of two replicates and we only included the genes detected in both replicates. Then we calculated the log10 transformed expression levels for following analysis.

Then we further filtered SVs with RNA-seq data and got 114,904 SV-gene pairs from 26 samples corresponding to 51,335 SVs and 20,318 genes. To match SVs with gene expression levels, we used the expression values of lead sample (the first sample detected corresponding SV) as target expression level. The RNA-seq data was generated from EBV-transformed lymphoblastoid cell lines (LCLs). Therefore, we used the SVEN (GM12878) to predicted the effects of these SVs on gene expression and compared the predicted values with observed values. Spearman correlation was used to evaluate the model performance. To further minimize the possibility of overestimating the performance due to the non-expressed genes, we removed all non-expressed genes and left 84,714 gene pairs, and this had negligible effects on the performance (Spearman correlation 0.915 on all pairs, 0.910 after removal).

**Estimation of effects of SVs in large-scale population and pathogenic SVs**

To better estimate the effects of SVs on gene expression in large-scale population, we applied SVEN on a SV dataset from 3,622 samples generated by Beyter *et al.* [16] using long-read sequencing. The dataset contains 133,886 SVs including 75,050 insertions, 55,649 deletions, and 3,187 unresolved SVs. We filtered SVs as the method described before and got 165,607 SV-gene pairs corresponding to 73,829 SVs and 22,235 genes. To estimate the effects of these SVs in population, we predicted the effects of these SVs in 53 GTEx tissues and calculated the max effect (absolute value) across all tissues.

Similarly, for pathogenic SVs, we used the SVs with clinical assertions from dbVar (nstd102, 20220829). We filtered SVs and only selected deletion (15,482) due to the lack of sequence information of other types of SVs. Here, we focused on high-confidence pathogenic deletions (clinical assertion is "Pathogenic") and we got 16,054 SV-gene pairs for 6,046 SVs and 9,118 genes. We also predicted the effects of these SVs in 53 GTEx tissues and calculated the max effect (absolute value) across all tissues.

**Experimental validation**

We selected deletions from mentioned SV datasets generated by Ebert et al. and Beyter et al. for experimental validation. To reflect the generalization ability of SVEN, we selected deletions (length < 1 kb) from four different cell lines and validated them in A375 cell line. Deletions are more likely to cause down-regulation of target gene, so we concentrated on the deletions predicted to cause up-regulation of genes. Finally, we selected top 5 deletions according to the effect on target gene in target cell line and validated these 5 deletions with CRSIPR experiment in A375 cell line.

Guide RNA pairs that were closest to the targeted SV boundaries were selected from UCSC genome browser CRISPR Targets track.

DNA fragments carrying guide RNA pairs (pgRNA) were generated by performing PCR on plasmid mU6-pgRNA-4.0 with primers pgRNA-F and pgRNA-R (**Supplementary Table 4**). These fragments were then examined with agarose gel electrophoresis, purified with the DNA Clean kit (Zymo), and assembled into the plasmid sgRNA-SV40-PURO using Golden Gate method. Success of these assemblies were confirmed by Sanger sequencings.

The pgRNA lentiviruses were produced by transfecting 70% confluent HEK293T cells with 1 μg sgRNA-SV40-PURO, 1 μg pCMVR8.74 (Addgene #22036), and 0.1 μg pVSV-G (Addgene #138479) in each well of a 6-well plate. The transfections were carried out using PEI (Proteintech). The lentiviruses were harvested by collecting supernatants of the 293T cell culture 72 hours post-transfection.

Monoclonal cell lines with constitutive Cas9 expressions were generated by transduction with lentivirus carrying a Cas9-2A-mCherry construct. These cells were then transduced with the target pgRNA lentiviruses at 70% confluency. 0.5 μg/mL puromycin (Invivogen) was added to the culture media 24 hours post-transduction until the cells reached 70% confluency again (~ 4 days). Cells were then cultured in fresh medium without puromycin for 24 hours before being collected. All cells were maintained in DMEM medium (HyClone) containing 10% fetal bovine serum (Gibco) and 1X penicillin-streptomycin-amphotericin B solution (Solarbio).

All nucleic acids from the cells were obtained by direct lysis on plates using the AllPure DNA/RNA kit (Magen), following the instructions.

We converted total RNA to cDNA by using HiScript III RT SuperMix for qPCR (Vazyme) following its instructions. We selected RPL41 as an internal reference gene as it has the most consistently high expression levels in the HPA database of 1055 cell lines. All qPCR reactions were performed in 96-well plates using LightCycler 480 (Roche). Each well contained 10 μL of ChamQ Universal SYBR qPCR Master Mix (Vazyme), 0.4 μL of 10 μM forward primer, 0.4 μL of reverse primer, and 9.2 μL of 1:25 diluted cDNA. Each set of measurements used 4-16 wells as technical replicates.
We performed melt curve analyses to ensure that the amplicons produced by the same pair of primers were specific and consistent each time. Each quantification cycle value was calculated using the Second Derivative Maximum method that came with the instrument, and the final value was the median between technical replicates. The specific details of all the above steps were carried out according to the manufacturers' instructions.
